# Astrocyte-derived extracellular vesicles as antigen-specific therapy for neuromyelitis optica spectrum disorder in the mouse model

**DOI:** 10.64898/2025.12.24.696382

**Authors:** Hamed Naziri, Gholamreza Azizi, Bogoljub Ciric, Guang-Xian Zhang, Abdolmohamad Rostami

## Abstract

Neuromyelitis optica spectrum disorder (NMOSD) is an autoimmune inflammatory disease of the central nervous system (CNS), characterized by Th17 cell responses and serum antibodies against the water channel aquaporin-4 (AQP4) on astrocytes. To avoid systemic immunosuppression by current therapies, an approach is to induce antigen (Ag)-specific tolerance by injecting an AQP4 epitope used to trigger the disease (e.g., AQP4_201-220_). The prerequisite for Ag-specific therapy is identification of the target Ag; however, dozens of epitopes of AQP4 and certain non-AQP4 astrocyte Ags have been identified as auto-Ags in NMOSD patients and in animal models. This uncertainty regarding relevant astrocyte Ags would hinder the translation of this and similar experimental strategies into Ag-specific therapy for NMOSD patients. In our study, we developed a therapeutic approach for an experimental NMOSD (eNMOSD) mouse model that relies on astrocyte-derived extracellular vesicles (AST-EVs), which theoretically contain all astrocyte Ags. Intravenous injection of AST-EVs mitigates disease progression in an AQP-4 Ag-dependent manner in the eNMOSD model with ongoing disease. AST-EVs suppressed inflammation by decreasing immune cell infiltration of the CNS, inducing T cell apoptosis, and increasing the frequency of regulatory T cells and IL-35-producing B cells. Furthermore, we defined that IFN-γ is crucial for successful i.v. tolerance induction by AST-EVs in eNMOSD. These novel findings represent a pioneering and considerable step toward a new therapeutic approach for NMOSD.

## Introduction

Neuromyelitis optica spectrum disorder (NMOSD) is an autoimmune disease of the central nervous system (CNS) that mainly affects the optic nerve and spinal cord, with minimal brain involvement (1). NMOSD is characterized by the production of autoantibodies against the aquaporin-4 (AQP4). Not all patients have these Abs, which are mainly expressed on astrocytes. The binding of these autoantibodies to AQP4 triggers immune-mediated cellular cytotoxicity, leading to direct astrocyte injury. This process subsequently results in secondary demyelination, oligodendrocyte damage, and neurodegeneration (2, 3). Current therapies for NMOSD are nonspecific and generally modulate the overall immune response. However, they remain, in most cases, only partially efficient (4). In addition, most immunotherapy focuses on B cells, the interleukin-6 receptor (IL-6R), and the complement system. These therapies have potentially serious side effects due to systemic immunosuppression, and they do not effectively eliminate autoreactive immune clones (5).

It is widely believed that autoimmune diseases develop due to a failure in peripheral immune tolerance for a specific self-antigen(s) (Ag) (6). Hence, an ideal therapy would re-establish tolerance for only the harmful immune responses targeting self-Ags, while leaving the rest of the immune system intact. The prerequisite for Ag-specific therapy is identification of the self-Ag. It is widely believed that NMOSD pathogenesis is driven by autoimmunity against AQP4 in astrocytes; however, the relevant self-Ag(s), i.e., various epitopes in AQP4 and non-AQP4 of astrocyte Ags, in NMOSD remain speculative, with the possibility that these Ags differ among patients (7–9). The complexity of auto-Ags and epitopes could impede the effectiveness of Ag-specific therapy using a single epitope. It would thus be best to develop Ag-specific therapies targeting all AQP4 epitopes without identifying relevant epitopes in each patient. In this regard, extracellular vesicles (EVs), which are double-lipid bilayer particles produced by various cell types that play an essential role in intercellular communication by transporting their contents to other cells, may be a solution (10). Our lab has demonstrated that EVs released from cultured primary oligodendrocytes contain all the relevant myelin Ags and suppress multiple experimental autoimmune encephalomyelitis (EAE) models of multiple sclerosis in an Ag-specific manner (11). These exciting findings provide a template for developing Ag-specific therapies targeting all auto-Ags of astrocytes in NMOSD, without the need to know the exact auto-Ag, and regardless of whether a patient is AQP4-Ab positive or negative.

A major challenge in research on NMOSD is the lack of an animal model that accurately simulates the disease (12, 13). Among the various NMOSD models, approaches encompass the administration of AQP4-IgG in EAE mice, passive transfer models involving direct injection of AQP4-IgG and human complement into the CNS, and adoptive transfer models of what? (14, 15). Notably, Serizawa et al. developed a model in which autoimmune NMOSD-like disease is induced through intradermal immunization with AQP4 peptide. This is the only model that replicates clinical manifestations of NMOSD (16).

EVs derived from astrocytes (AST-EVs) contain multiple proteins with both neurotrophic and neuroprotective features (17, 18). AST-EVs enhance the survival rate and function of neurons exposed to oxidative stress and are involved in the remyelination and CNS repair by modulating microglia (19). Here, we investigated the therapeutic potential of AST-EVs in experimental NMOSD (eNMOSD) mouse model. We established that the intravenous (i.v.) administration of AST-EVs effectively inhibits disease progression in an eNMOSD mouse model in an Ag-specific manner. These novel findings represent a pioneering step toward a new therapeutic approach for NMOSD.

## Material and Methods

### Mice

Adult female C57BL/6 mice (8-10 weeks) were purchased from The Jackson Laboratory (Bar Harbor, ME, USA). Mice were housed in clean cages, with a maximum of five mice per cage in a controlled environment with 12 h/12 h of light/dark cycles and unlimited access to food throughout the experimental procedures. All procedures were taken to minimize the pain and distress of the animals. All experiments were conducted with prior Institutional Animal Care and Use Committee approval.

### eNMOSD induction and i.v. tolerance

Mice were injected intradermally at 15 sites from the base of the tail to the back, with 100 µL of an emulsion containing 400 µg of AQP4_201-220_ peptide (Genscript, NJ, USA) and an equal volume of Complete Freund’s adjuvant supplemented with 10 mg/mL of heat-killed *Mycobacterium tuberculosis* H37Ra. Additionally, mice were i.v. injected with 500 ng of pertussis toxin on Day 0 and intraperitoneally on Day 2 (16). Clinical assessment of eNMOSD was conducted according to the following scale: 0 = healthy; 1 = limp tail; 2 = ataxia and/or paresis of hindlimbs; 3 = paralysis of hindlimbs and/or paresis of forelimbs; 4 = tetraparalysis; and 5 = moribund or death. I.v. tolerance was induced as previously described (20). Briefly, after disease onset, each mouse received either 100 μg AQP4_201-220_ in PBS or at least 10^10^ AST-EVs in PBS, every third day, 2 times in total. Control mice received PBS or 10^10^ HEK-EVs.

### HEK cells

HEK293 cells were purchased from the American Type Culture Collection (CRL-1573, ATCC, United States). HEK cells were cultured in Dulbecco Modified Eagle’s Medium (DMEM, Gibco) supplemented with 10% EV-depleted fetal bovine serum (Gibco™; Cat # A2720801), penicillin, streptomycin (100 U/ml), and 2 mM L-glutamine. All cells were maintained at 37°C in 5% CO2.

### GLAST (ASCA-1) isolation and astrocyte culture

Whole brains were harvested from 3-5-day-old C57BL/6 mouse pups, manually dissociated, and a neural dissociation kit (Miltenyi; Cat # 130-092-628) was used for enzymatic digestion. The digestion was terminated with DMEM (Gibco; Cat # 11965092), supplemented with 10% EV-depleted FBS, and centrifuged at 1200 rpm for 5 min. The tissue was then homogenized by passing through an 18-gauge needle and filtering through a 70 mm cell strainer to remove debris. A positive selection with magnetic beads separation kit (Miltenyi; Cat # 130-095-826) was utilized to isolate GLAST^+^ cells (ASCA-1^+^). The cells were then cultured in DMEM containing 10% FBS at 37°C/5% CO_2_ for 6–10 days until confluence and maturity. Medium was replaced every 3 days.

### EV purification

The mature astrocyte culture supernatants were collected for EV isolation every 2–3 days using a differential centrifugation (21). The supernatants were centrifuged at 300 g for 10 min to remove live cells and debris. Then, to remove dead cells, the resulting supernatants were further centrifuged at 2,000 g for 10 min, followed by 10,000 g for 35 min at 4°C to remove large vesicles and apoptotic bodies. For the final step, supernatants were centrifuged at 110,000 g for 70 min at 4°C. Depending on their intended application, the pellets were resuspended in either lysis buffer supplemented with protease inhibitors, 0.1 μm-filtered PBS, or fixative.

### Nanoparticle Tracking Analysis (NTA)

Diluted EVs (1:2,000 in PBS) were measured using the ZetaView® x30 instrument (Particle Metrix GmbH). Particle visualization, particle number concentration, and particle size distribution were determined by NTA.

### Histological evaluation

On day 23 post immunization (d.p.i.), mice were thoroughly perfused with ice-cold PBS (pH 7.4, 100 mL per mouse) through the left cardiac ventricle in each group. The spinal cord and optic nerve were dissected and then fixed in 4% paraformaldehyde (PFA) in PBS overnight. The following day, tissues were transferred to 70% ethanol and incubated overnight. Tissue processing was conducted on the fixed tissue, and then it was embedded in paraffin to create a paraffin block. Ten-micrometer sections were cut and placed on slides. for staining, deparaffinization was conducted by xylene and then decreasing graded concentrations of alcohol for 1 minute each. Sections were stained by hematoxylin for 2 minutes, washed, and then stained by eosin for 1 minute. LFB staining was performed on a deparaffinized slide with LFB solution (Sigma-Aldrich) at 56°C overnight.

### Preparation of CNS-infiltrating mononuclear cells (MNCs) and splenocytes

After extensive perfusion, brains and spinal cords were mechanically dissociated using scissors, followed by enzymatic digestion with 700 µl/mL Liberase TL (Sigma Aldrich, USA) at 37°C for 30 minutes to isolate MNCs. cells were suspended in a one layer of 40% Percoll (GE Healthcare, Little Chalfont, U.K.) and spun at 500 × *g* for 30 min at room temperature. Spleens were disrupted using a 70 µm cell strainer before being treated with red blood cell lysis buffer (Miltenyi) for ∼2 minutes to obtain splenocytes. Cold PBS was utilized to wash harvested cells for flow cytometry analysis.

### Flow cytometry

The surface-marker staining was performed by incubating cells with fluorochrome-conjugated Abs: CD45-Alexa fluor 700, CD4-FITC, CD8-PE, CD11c- Percp-cy5.5, CD11b-DAPI, Ly6c-APC, Ly6G-PE/Cy7, PD-1-PE, CD19-FITC and CD11C-PE/Dazzle™ 594 (Biolegend and BD Biosciences) in two different panels at the recommended dilution or FMO control Abs for 30 min on ice. Intracellular staining, cells were stimulated PMA (50 ng/ml; Sigma-Aldrich), Ionomycin (500 ng/ml; Sigma-Aldrich), and GolgiPlug (1 μg/ml; BD Biosciences) at 37°C for 5 h. Cell permeabilization was performed using Fix and Perm Medium B (Thermo Fisher). Antibodies targeting intracellular antigens were added in a total volume of 200 μL (Fix and Perm Medium B from Thermo Fisher and PBS/3%FBS at a 1:1 ratio) and incubated for 1 h or overnight. Abs against intracellular antigens included: IL-17-BV786, IFN-g-BV711, GM-CSF-PE/Dazzle™ 594, IL-10-PE/Cy7, Caspase-3-BV650/Cy7, EBI3-APC, IL-10-DAPI, IL-27(P28)-PE/Cy7, and IL-35(P35)-PE, from Biolegend and BD Biosciences in two different panels. Dead cells were excluded using LIVE/DEAD™ Fixable Yellow Dead Cell Stain Kit (ThermoFisher; Cat # L34959). Data were acquired on a FACSAria Fusion (BD Biosciences) and analyzed by FlowJo software (TreeStar).

### Western blot

EVs were lysed by RIPA buffer (Thermo Scientific, Cat # 89900) supplemented with 1× proteinase inhibitor cocktail (Thermo Scientific, Cat # 87786). Protein concentrations were determined with BCA (Micro BCA, Pierce). Totally, 5-10 μg of protein from lysed EVs were diluted with Laemmli buffer and loaded onto a 4–12% gradient SDS-PAGE. The following primary antibodies were used: Rabbit anti-AQP4 (R&D Systems, Cat # NBP1-87679), mouse anti-mouse flotillin1 (BD Bioscience, Cat # 610821), goat anti-Tsg101 (Millipore), and mouse anti-GAPDH (Millipore, Cat # CB1001). Incubation with secondary antibodies was performed using anti-goat, anti-rabbit, and anti-mouse IgG-horseradish peroxidase (HRP) antibodies (Thermo Scientific).

### ELISA

Splenocytes were cultured at a density of 2×10^5^ cells per well in a flat-bottom 96-well plate containing IMDM supplemented with 10% heat-inactivated fetal calf serum, 2 mM L-glutamine, 100 U/mL penicillin, 100 μg/mL streptomycin, and 50 μM β-mercaptoethanol. Splenocytes were stimulated with 20 µg of AQP4_201-220_ for three days. Then the supernatants from cell cultures were collected and stored at −20 ◦C until use. The concentrations of IL-17, GM-CSF, IFN-γ, TNF, and IL-10 in culture supernatants were determined using an enzyme-linked immunosorbent assay (ELISA) using commercial kits (R&D Systems, Minneapolis, MN, USA).

### AQP4_201-220_-Specific autoantibodies quantification in sera of mice with EAE

Biotinilation of AQP4_201-220_ was performed using EZ-Link Sulfo-NHSBiotin (Thermo Fisher Scientific, cat# A39256). Then, streptavidin-coated plates (Thermo Fisher Scientific, cat#15120) were coated with biotinylated peptide for 1 h at room temperature. Following 3 times of washing, the plate was subsequently incubated with diluted sera (1/50) for 1 h at room temperature. After washing, HRP-conjugated goat anti-mouse IgG H&L (Abcam cat# ab97023) was added and incubated for 1 h at room temperature. For the colorimetric reaction, TMB (Thermo Fisher Scientific, cat#34028) was added to the wells, and the reaction was terminated by adding sulfuric acid.

#### Statistical analysis

Statistical analysis was performed using GraphPad Prism 8 software. Statistics were evaluated using an unpaired, two-tailed Student’s t-test between two groups and a Kruskal-Wallis test between three or more groups for clinical score analysis. *P* ≤ 0.05 was considered significant. Data represent means ± SEM.

## Results

### AQP4_201-220_ peptide suppresses disease progression in eNMOSD

Induction of Ag-specific tolerance by i.v. injection of an encephalitogenic peptide in EAE has been extensively studied, including in our laboratory (11, 22). To evaluate whether the AQP4_201–220_ peptide can suppress the progression of ongoing eNMOSD, we immunized C57BL/6 mice with AQP4_201–220_ peptide emulsified in CFA with pertussis toxin, as described (16). We then examined the effect of AQP4_201–220_ administered via i.v. injection at the onset of clinical disease on days 12 and 15 p.i. (50 μg/dose). Administration of the peptide resulted in a profound and sustained suppression of clinical disease. (**Fig. S1A and B).** The mice exhibited significantly reduced numbers of infiltrated cells **(Fig. S1C)** and CD45^high^ cells (leukocytes; **Fig. S1D**) in the CNS compared with control mice that were treated with PBS. Analysis of CNS immune cell populations, as gated in **Fig. S3**, revealed a significant decrease in the absolute numbers of isolated CD4⁺ T cells, B cells, NK cells, and CD11b⁺Ly6C^high^ (monocytes) in the AQP4_201–220_ peptide-treated group compared to the PBS-treated group; however, their relative frequencies did not differ significantly between the groups **(Fig. S1E)**. Non-activated microglia (CD45^lo^CD11b⁺) exhibited a smaller difference in absolute numbers than other cell types.

The frequencies and absolute numbers of proinflammatory cytokine–producing CD4⁺ T cells were investigated. Absolute numbers of IL-17A, IFN-γ, and GM-CSF-producing CD4⁺ T cells were markedly higher in mice treated with PBS compared to those treated with AQP4_201-220_; however, there was no significant difference between groups in their frequencies (**Fig. S2A and C**). While the frequency of Foxp3⁺ and IL-10⁺ CD4⁺ T cells was slightly higher in AQP4_201-220_ -treated mice, these differences were not statistically significant. Again, if it is not significant, then it is not different, and writing about it like “slightly higher” does not make sense. The absolute numbers of Foxp3⁺ and IL-10⁺ CD4⁺ T cells were reduced in the AQP4_201-220_-treated mice compared to PBS-treated mice. (**Fig. S2B and C**). In summary, these data show that i.v. injections of AQP4_201-220_ peptide suppress disease in the eNMOSD mouse model.

### EVs secreted by astrocytes contain AQP4

To determine whether AST-EVs carry AQP4, we first established a highly enriched population of primary astrocytes and characterized the EVs they secrete. Astrocytes isolated from C57BL/6 neonatal brains showed high purity, with flow cytometric analysis demonstrating that approximately 93% of the cells were positive for the astrocyte marker GLAST (**Fig. 1A**). Astrocytes were further confirmed by robust expression of glial fibrillary acidic protein (GFAP) as assessed by immunofluorescence staining (**Fig. 1B**) Astrocytes produced large quantities of EVs (2.5 × 10^11^/mL) with an average diameter of 160 nm, as determined by nanoparticle tracking analysis (NTA) (**Fig. 1C**). Western blot analysis showed that AST-EVs, but not control HEK-EVs, contain AQP4. Principal EV markers (e.g., Flotillin-1, TSG101), according to minimal information for studies of extracellular vesicles (MISEV) guidelines, were also present in AST-EVs (**Fig. 1D**). Representative transmission electron microscopy (TEM) images showed the spherical shape and the size of mouse EVs (**Fig. 1E**). The presence of AQP4 in purified AST-EVs was confirmed by (**Fig. 1F**). Overall, these data show that astrocytes release EVs containing AQP4, making them a promising option for tolerance induction in eNMOSD.

**Fig. 1.**
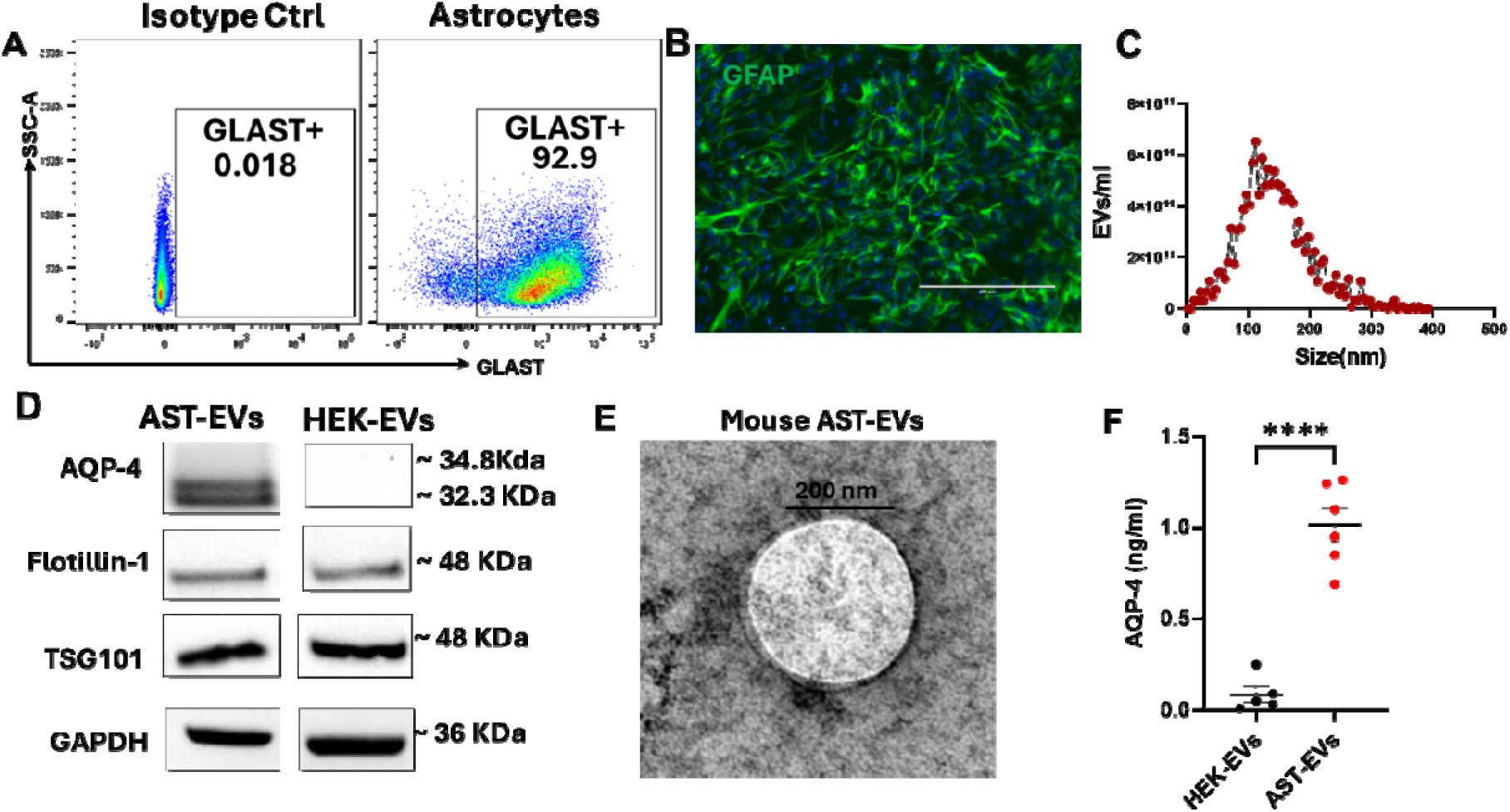
Characterization of astrocytes and AST-EVs. **(A)** Flowcytometry analyses for GLAST^+^ in astrocytes isolated from the CNS of 5–7-day-old mouse pups. **(B)** Representative immunofluorescence of GFAP (green) for astrocytes, and nuclei (blue). Scale bar, 20 nm. EVs were purified from the supernatants of mouse astrocyte culture and characterized for size and protein content. **(C)** Size profile of AST-EVs determined by NTA. **(D)** Representative Western blot for AQP4, FLOT-1, TSG101 and GAPDH in AST-EVs and HEK-EVs pellets. (**E**) Representative TEM image shows purified mouse AST-EVs (scale bar: 200 nm; HV = 80.0 kiva). **(F)** Concentrations of AQP4 proteins in AST-EVs lysate were determined by ELISA. HEK-293 cell (HEK)-derived EVs was used for comparison. Values are mean ± SD of 5-6 preparations/group. Unpaired t-test; ****p < 0.0005.

### AST-EV treatment alleviates clinical severity and CNS inflammatory demyelination in eNMOSD mice

To assess the therapeutic efficacy of AST-EVs, eNMOSD was induced as described in Fig. 1, followed by i.v. administration of AST-EVs or HEK-EVs (as control) at a dose of 10^10^ particles per injection on days 13 and 16 p.i. (at disease onset; indicated by red arrows, **Fig. 2A**). AST-EV-treated mice showed a significantly less severe disease course and bodyweight loss compared to the HEK-EV-treated group (**Fig. 2A, B**). Significantly fewer inflammatory cell infiltrates and reduced demyelination were observed in AST-EV-treated mice than in HEK-EV-treated mice, as determined by H&E and LFB staining of the spinal cord (**Fig. 2C-E**) and optic nerve (**Fig. 2F-H**). Overall, the effect of AST-EVs was similar to that of AQP4_201–220._ These data reveal that i.v. injections of AST-EVs successfully attenuate ongoing clinical disease in eNMOSD mice.

**Fig. 2.**
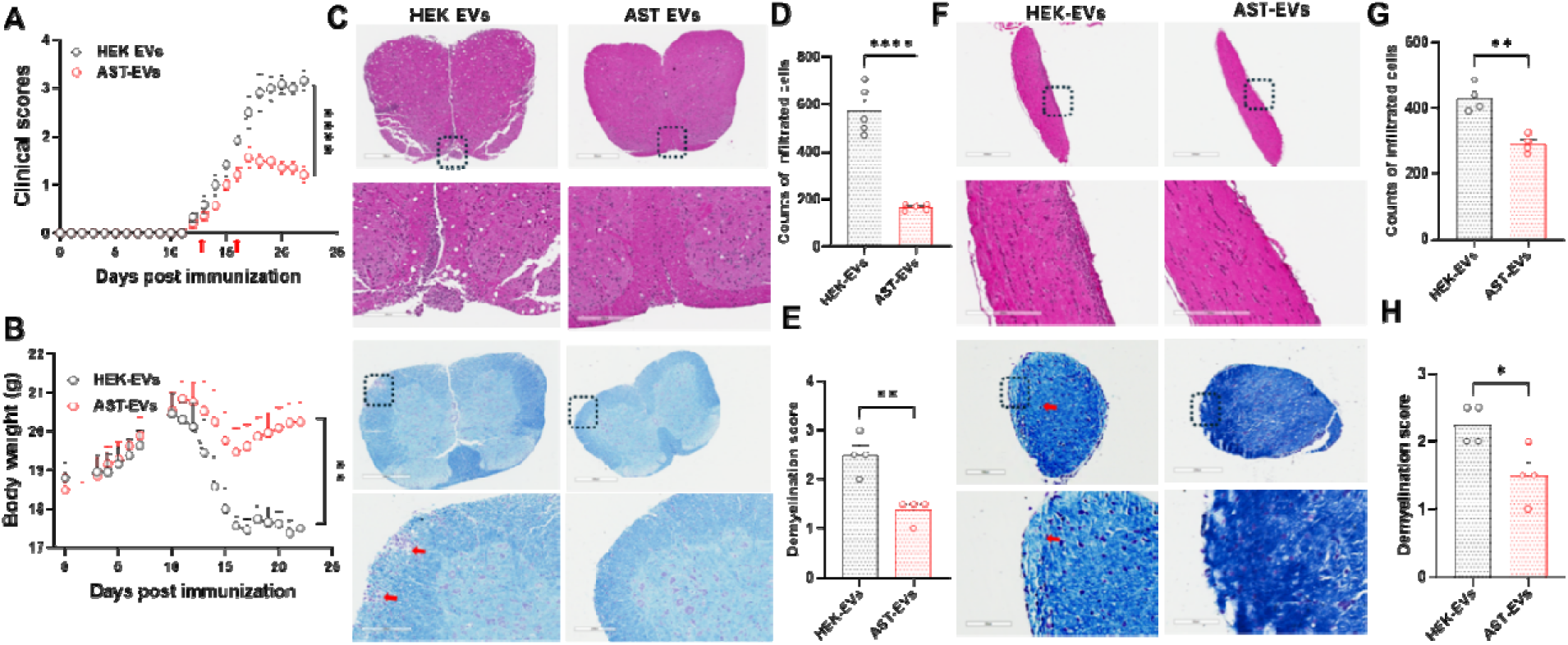
AST-EVs/i.v. ameliorate clinical disease in eNMOSD. Approximately 10^10^ of AST-EVs (prepared from astrocytes of C57BL/6 mice) or HEK-EVs were i.v. injected (red arrows) in AQP4_201-220_-induced eNMOSD mice. Injections were given i.v. at disease onset (days 13 and 16 p.i.). Clinical course **(A)** and body weight **(B)** were recorded. **(C-H)** H&E and LFB staining of paraffin sections of spinal cord **(C-E)** and optic nerves **(F-H)** isolated from HEK-EV- and AST-EV-treated NMOSD mice on day 23 p.i. Values are mean ± SD of 5-6 mice/group. *P < 0.05, **P < 0.001, and **** P < 0.0001. One experiment representative of the two experiments is shown.

AST-EV treatment significantly reduced the total number of cells isolated from the CNS, which was due to reduced numbers of CD45^hi^ cells infiltrated from the periphery (**Fig. 3A**). A marked decrease in the numbers of CD4⁺ and CD8⁺ T cells was observed in AST-EV-treated mice, although their relative percentages remained unchanged (**Fig. 3B**). While percentages of CD11b⁺Ly6G⁺ neutrophils among CD45^hi^ cells, CD11b⁺CD45^lo^ microglia, CD11b⁺Ly6C^hi^ monocytes and CD11c⁺ dendritic cells were similar between AST-EV- and HEK-EV-treated mice, their absolute numbers were significantly reduced in the AST-EV group (**Fig. 3C-E**). These findings demonstrate that AST-EVs can be utilized for i.v. tolerance induction in ongoing eNMOSD.

**Fig. 3.**
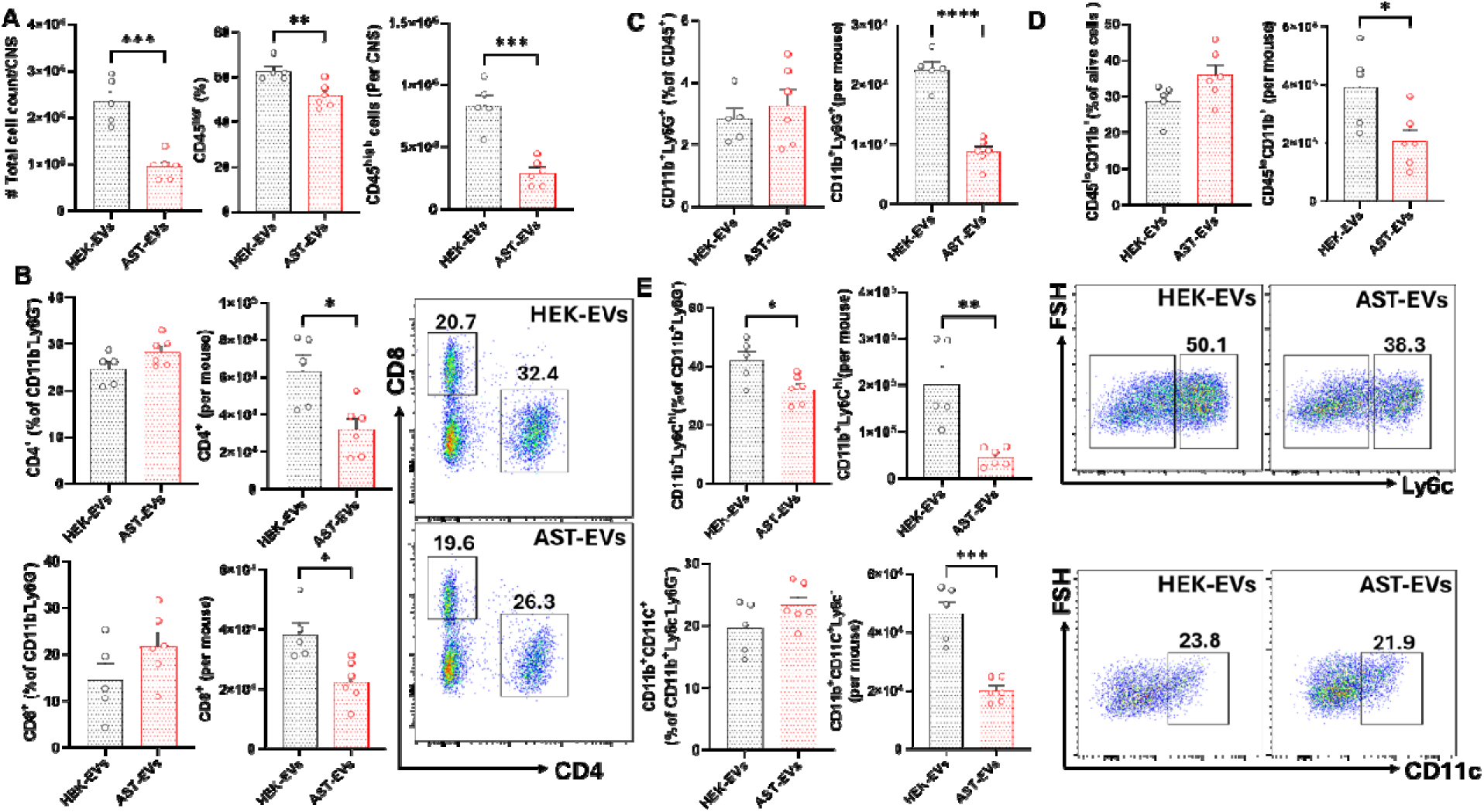
AST-EV treatment decreases immune cell infiltration into the CNS. Mice shown in Fig. 2 were sacrificed on day 23 p.i., and mononuclear cells (MNCs) in the CNS were isolated. The numbers of CD45^hi^ infiltrating leukocytes were determined by flow cytometry (A). The frequencies and absolute numbers of CD4^+^ T cells, CD8^+^ T cells, neutrophils, microglia, monocytes, and DCs are also shown (B-E). Results are expressed as the mean + SEM, with n≥ 5 per group from two independent experiments. Data were analyzed by Student’s t-test; *P < 0.05; **P < 0.01; ***P < 0.001; ****P < 0.0001.

### AST-EV treatment modulates cytokine production and suppresses Ag-specific proliferation of splenocytes from eNMOSD mice

To investigate the production of inflammatory cytokines by CNS-infiltrating CD4^+^ T cells, the frequencies and absolute numbers of IFN-γ+, GM-CSF+, IL-17+, and TNF+ cells were determined by flow cytometry. Although we did not find difference in the frequencies between the two groups, the absolute number of cells producing inflammatory cytokines was higher in HEK-EVs-treated mice (**Fig. 4A**). AST-EVs/i.v. injection increased the frequency of IL-10^+^CD4^+^ T cells (**Fig. 4B**) and induced programmed death-1 (PD-1) expression on CD4^+^ T cells (**Fig. 4C**). These data show that AST-EVs can induce an anti-inflammatory response and increase apoptosis in CD4^+^ T cells.

**Fig. 4.**
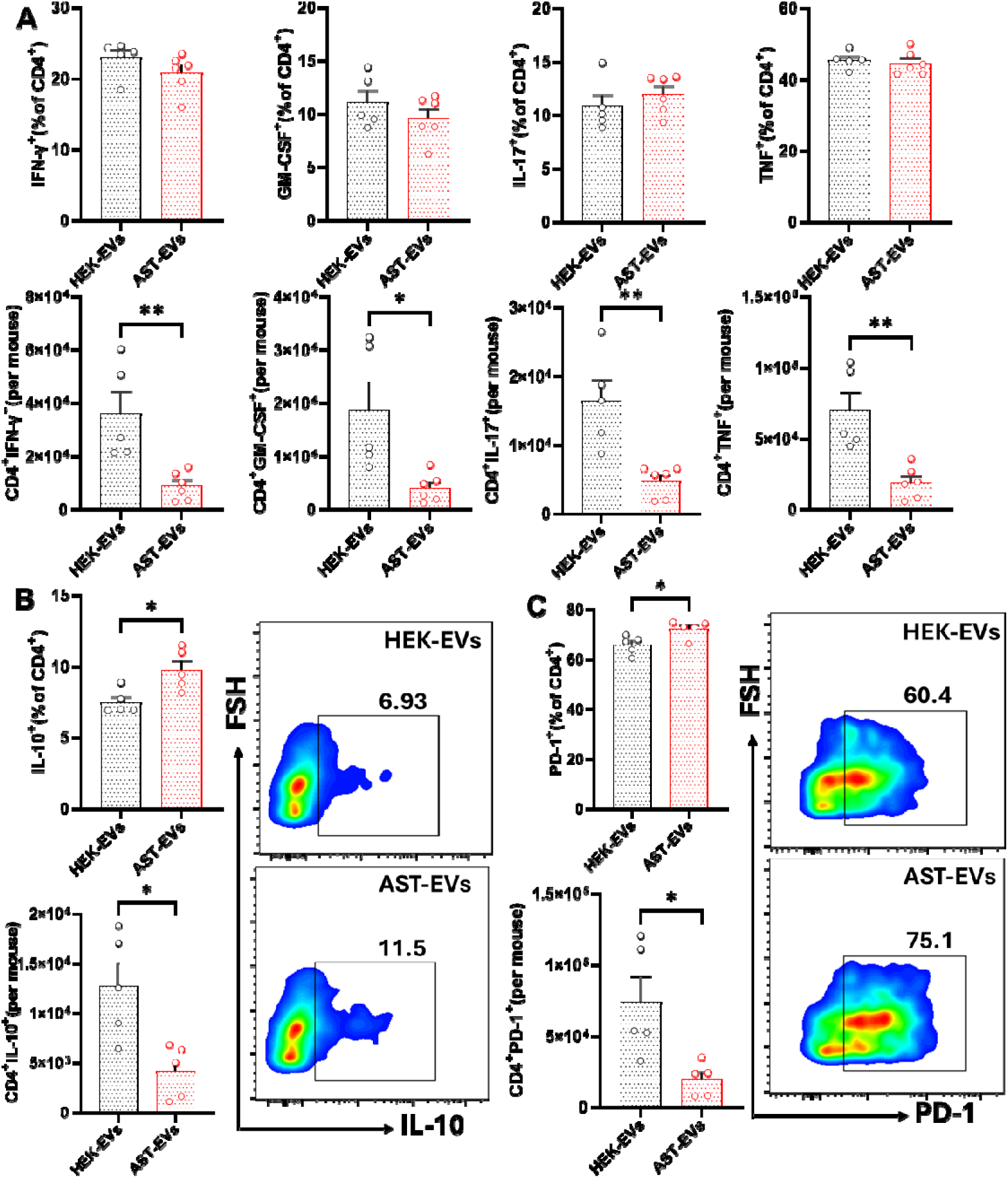
Frequency and absolute number of cytokine-producing T cells in the HEK-EV- and AST-EV-treated mice. (**A**). Mice shown in Fig. 2 were sacrificed on day 23 p.i., and MNCs in the CNS were isolated. Higher absolute numbers of cytokine-producing CD4^+^ T cells were observed in the HEK-EV-treated during eNMOSD. (**B**) IL-10^+^CD4^+^ T cell frequencies were significantly higher in AST-EV-treated mice. (**C**) PD-1 expression frequencies and absolute numbers in CD4^+^ T cells. Data were analyzed by Student’s t-test; *P < 0.05; **P < 0.01.

We then determined the effects of AST-EVs on the peripheral immune system. Splenocytes from eNMOSD mice treated with either AST-EVs or HEK-EVs were isolated and stimulated by AQP4_201-220_. Cytokine levels in the culture supernatants were measured using ELISA after 72 hours of activation. Splenocytes of AST-EV-treated mice produced lower levels of IL-17 and GM-CSF, but higher levels of IL-10 compared to the HEK-EV group (**Fig. 5A**), indicating an increased regulatory potential. There were no differences in IFN-γ or TNF levels. To further characterize AQP4_201-220-_specific CD4^+^ T cell response in AST-EV-treated mice, we tested proliferative response by XTT assay. Splenocytes of AST-EV-treated mice proliferated less than those of control mice, whereas there was no difference for anti-CD3/anti-CD28 Ab-stimulated Ag-non-specific proliferative response (**Fig. 5B**). These results support the possibility that AST-EV treatment is Ag-specific without systemic immunosuppression.

**Fig. 5.**
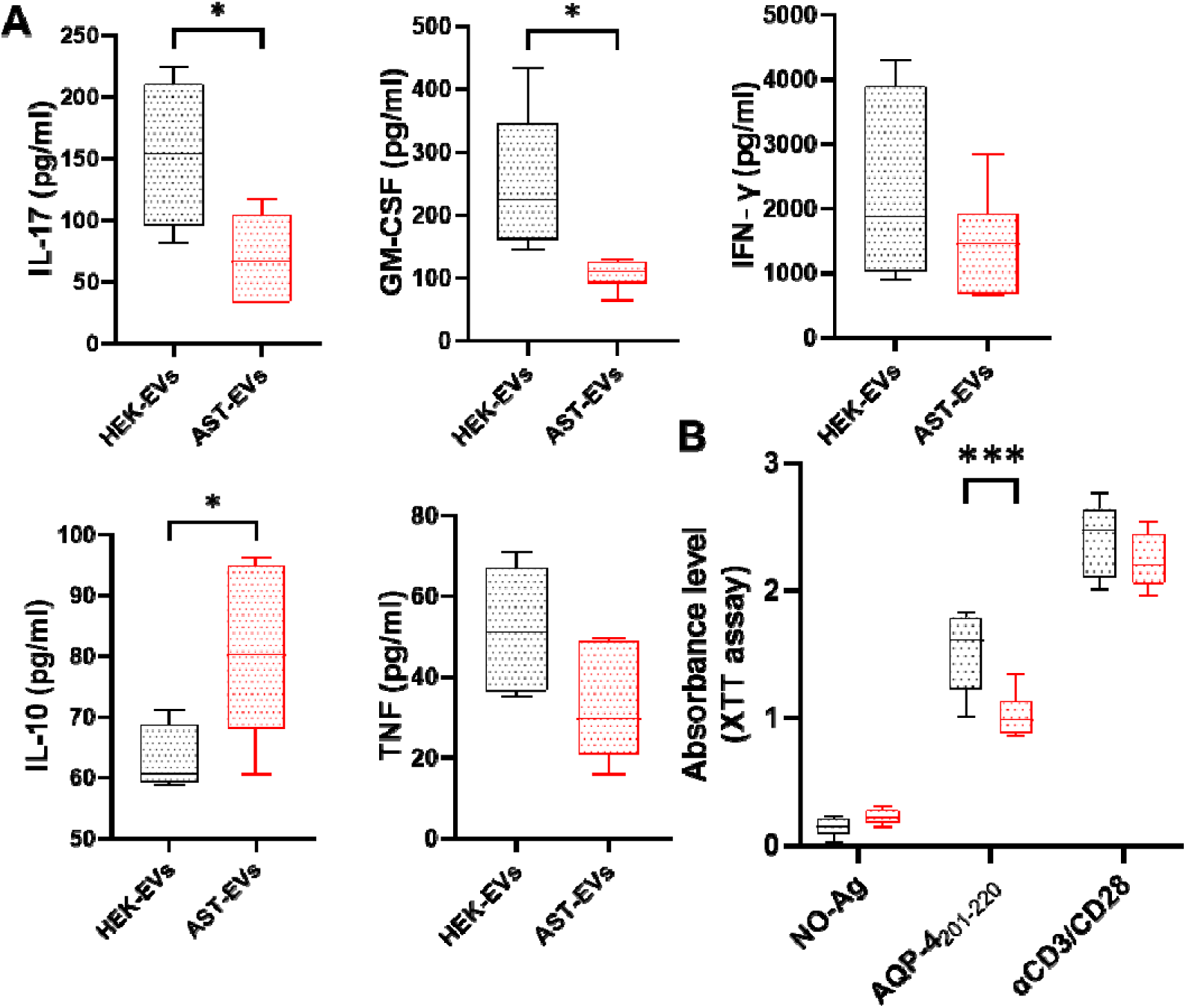
AST-EVs inhibit inflammatory cytokines and Ag-specific T cell responses. For cytokine measurement and proliferation assay, splenocytes from the mice shown in Fig. 2 were seeded (3×10^5^ cells per well) in a 96-well plate and activated with AQP4_201-220_. (**A**) Cytokine concentrations in the supernatants of splenocyte cultures stimulated with AQP4_201-220_ in HEK-EVs or AST-EVs were determined by ELISA. (**B**) Splenocytes from mice were cultured for 4 days with either no antigen (No-Ag), AQP4_201-220_ peptide, or anti-CD3/CD28 antibodies. Absorbance levels, suggestive of cell proliferation, were measured at 450 nm using the XTT assay. Data represent mean ± SEM (n=5 each group). Data were analyzed by Student’s t-test; *P < 0.05; **P < 0.01; ***P < 0.001.

### AST-EV treatment promotes IL-35-producing Breg cells, reduces the ABC subset in the CNS, but does not impact serum levels of anti-AQP4 Abs

To determine the impact of AST-EVs on CNS-infiltrating B cells, we evaluated CD19⁺ B cells in eNMOSD mice by flow cytometry. AST-EV-treated mice showed a significant decrease in the frequency of total CD19⁺ B cells (**Fig. 6A**). A significantly higher frequency of B cells expressed IL-35 (EBI3⁺P35⁺), suggestive of regulatory B cell (Breg) activity (23). The expression of IL-27 (EBI3^+^P28^+^) and IL-6 by B cells did not differ between the two groups. (**Fig. 6B**). The proportion of CD11c⁺ cells among CD19⁺ cells, a group of autoimmune-associated B cells (ABCs) (24), was reduced in AST-EV-treated mice (**Fig. 6C**). No significant difference was observed in serum anti-AQP4 IgG levels between these two treatment groups (**Fig. 6D**), suggesting that AST-EVs modulate B cell function rather than their Ab production.

**Fig. 6.**
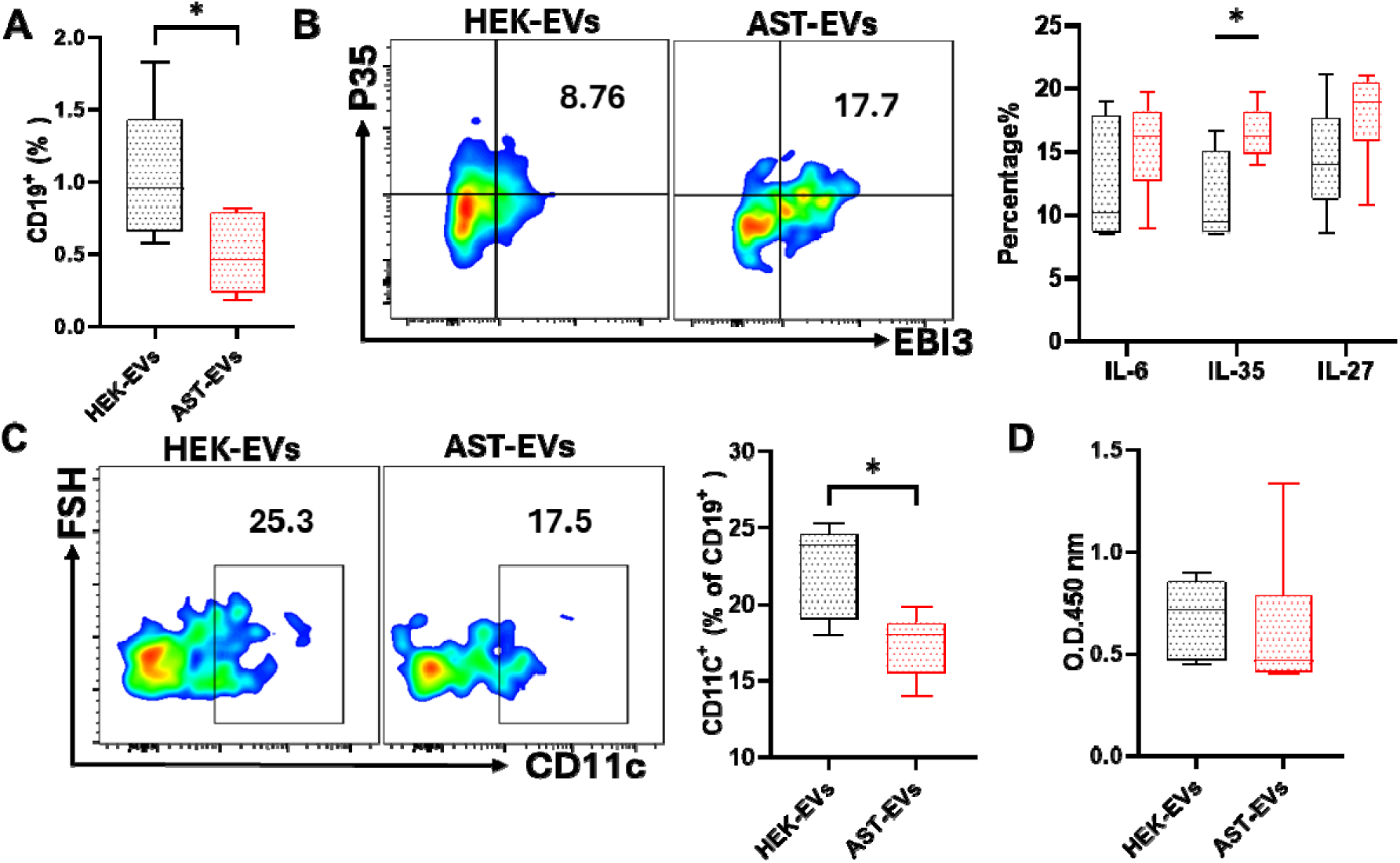
The frequency and cytokine production of B cells in the CNS and serum anti-AQP4 Ab levels. Mice shown in Fig. 2 were sacrificed on day 23 p.i., and MNCs isolated from the CNS were analyzed for CD19⁺ B cells (A), their production of IL-35 (EBI3⁺p35⁺), IL-27 (EBI3⁺p28⁺), and IL-6 (B), and expression of CD11c⁺ (C). Representative flow cytometry plots and quantification are shown. (D) Anti-AQP4 auto-Abs were measured by ELISA in the sera of eNMOSD mice treated with AST-EVs and HEK-EVs. Optical density (OD) at 450 nm is shown. Data were analyzed by Student’s t-test (n=5 each group); *P < 0.05.

### AST-EVs enhance the apoptosis of CD4⁺ T cells in eNMOSD mice

To determine if AST-EV treatment induced apoptosis of T and B cells, active Caspase-3 expression was measured in CD4^+^ T cells and CD19^+^ B cells from the CNS and spleen of eNMOSD mice following the treatment. Mice treated with AST-EV had significantly greater proportion of Caspase-3^+^CD4^+^ T cells in both the CNS and spleen compared to mice treated with HEK-EV (**Fig. 7A**). However, there was no difference in the proportion of Caspase3⁺CD19⁺ B cells between the treatment groups in either tissue (**Fig. 7B**). Moreover, the PD-1 expression on CD4^+^ T cells has been evaluated to determine their potential role in regulating T cell activation and exhaustion upon AST-EV treatment. The treatment increased the percentage of PD-1^+^CD4^+^ T cells in the CNS, but not in the spleen, compared to HEK-EV treatment (**Fig. 7C**). These results show that apoptosis of CD4^+^ T cells may be a mechanism underlying AST-EV-induced tolerance in eNMOSD.

**Fig. 7.**
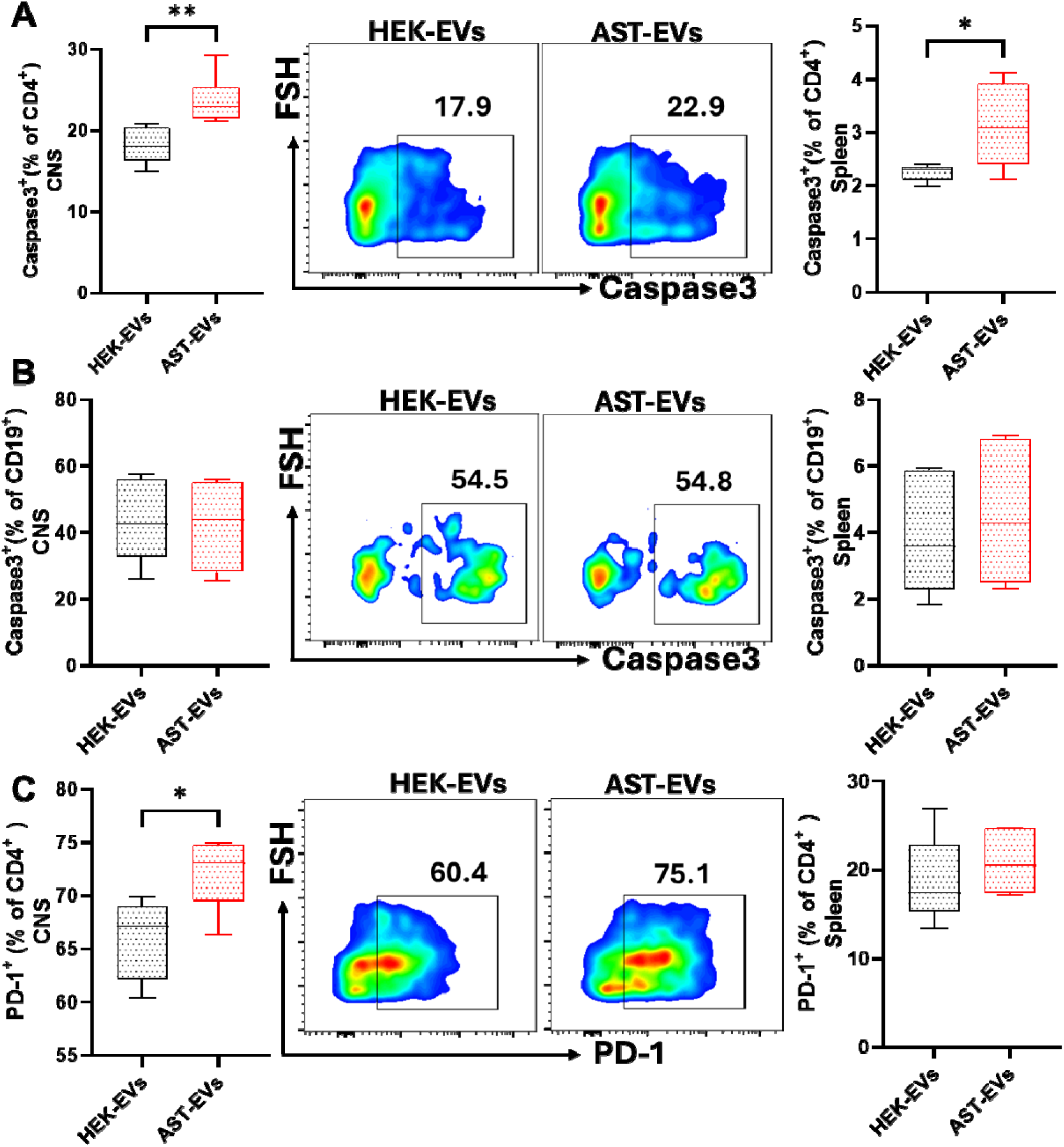
Apoptosis analysis of T and B cells in the CNS and spleen following treatment with HEK-EVs or AST-EVs. Mice shown in Fig. 2 were sacrificed on day 23 p.i., and MNCs isolated from the CNS and spleen were analyzed. (**A**) Frequency of apoptotic CD4⁺ T cells (Caspase-3⁺) in the CNS (left) and spleen (right) assessed by flow cytometry. (**B**) Frequency of apoptotic CD19⁺ B cells (Caspase-3⁺) in the CNS (left) and spleen (right). (**C**) The frequency of PD-1 expression in CD4⁺ T cells in the CNS (left) and spleen (right) was assessed by flow cytometry. Results are expressed as the mean ± SEM with n≥ 5 per group from 2 independent experiments. Data were analyzed by Student’s t-test (n=5 each group); *P < 0.05, and **P < 0.01.

### IFN-γ is essential for i.v. tolerance induction by AST-EVs in eNMOSD

We have described that IFN-γ signaling plays an important role in myelin Ag peptide-induced tolerance in EAE (25). Here, we tested if this is also the case for EV-induced tolerance in eNMOSD. While treatment with AST-EVs significantly suppressed eNMOSD, this effect was largely precluded by injection of neutralizing anti-IFN-γ mAb 24 h before AST-EVs injection (**Fig. 8A and B**). Mice treated with anti-IFN-γ mAb had significantly elevated numbers of infiltrated cells and CD45^high^ cells (leukocytes) in the CNS compared with control mice treated with isotype mAbs (**Fig. 8C**). The CNS of anti-IFN-γ + AST-EV-treated and control mice showed no difference in the mean frequency of isolated CD4^+^ T cells, CD8^+^ T cells, CD11b^+^Ly6G^+^ (neutrophils), CD45^lo^CD11b^+^ (microglia), CD11b^+^Ly6C^high^ (monocytes) and CD11b^+^CD11c^+^ (DCs) at the peak of eNMOSD (23 days p.i.; **Fig. 8D**). However, the cell number isolated from the CNS of anti-IFN-γ+ AST-EVs treated mice was lower at the peak of the disease compared to control mice (**Fig. 8E**).

**Fig. 8.**
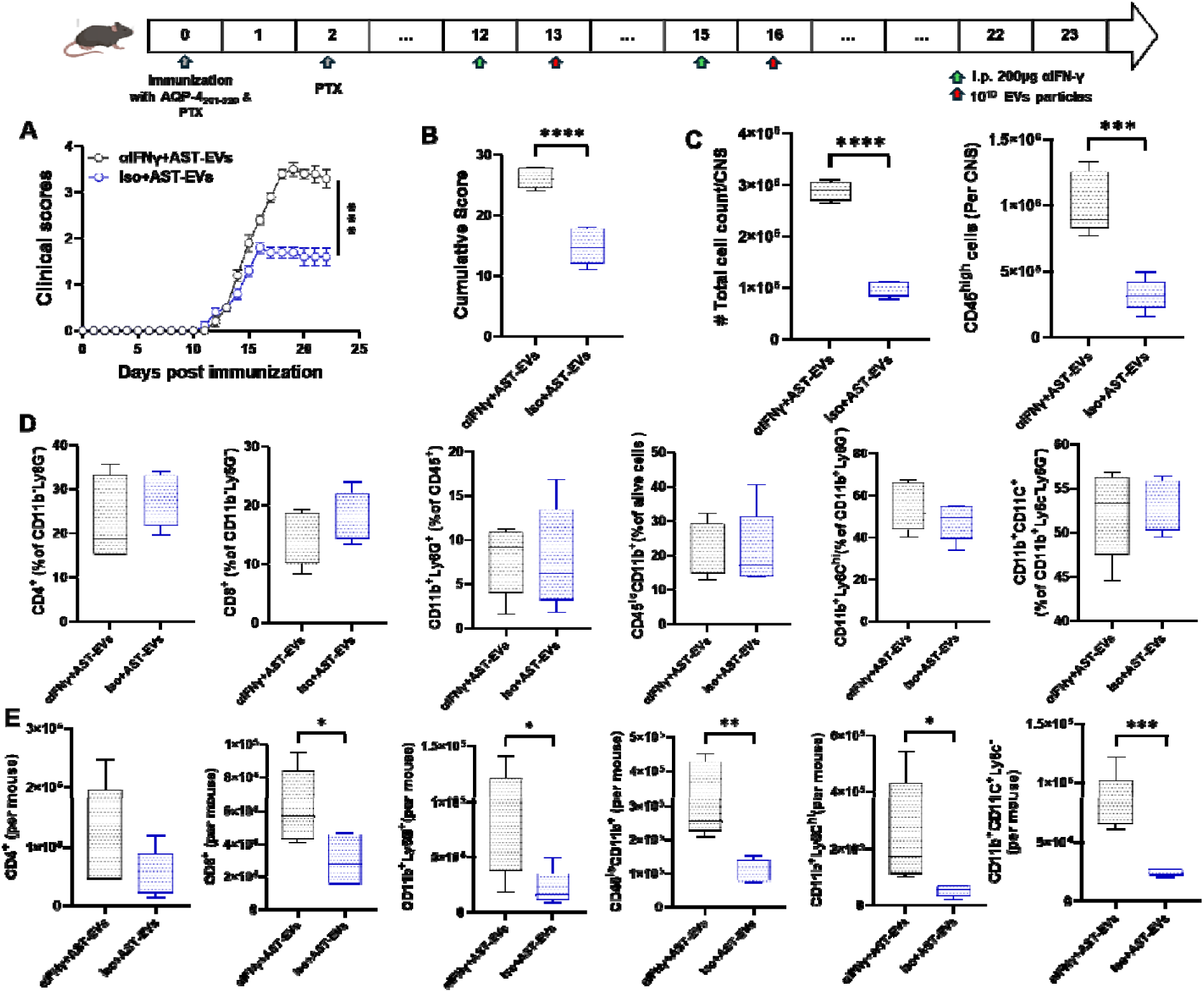
IFN-γ is critical for i.v. tolerance induction in eNMOSD by AST-EVs. (A) Mice with eNMOSD (n = 5 per group in each experiment) were treated via intraperitoneal (i.p.) injection with a blocking anti-IFN-γ MAb (200 μg per mouse per injection; clone XMG1.2) or an isotype control MAb on days 12 and 15 p.i. AST-EVs were administered i.v. on days 13 and 16 p.i. (B) Cumulative scores of clinical severity. (C) Mice were sacrificed on 23 d.p.i., and the numbers of CD45^high^ leukocytes in their CNS were determined by flow cytometry. (D) The frequencies and absolute numbers of CD4^+^ T cells, CD8^+^ T cells, CD11b^+^Ly6G^+^ (neutrophils), CD45^lo^CD11b^+^ (microglia), CD11b^+^Ly6C^hi^ (monocytes), and CD11b^+^CD11c^+^ (DCs) are also shown. Results are expressed as the mean + SEM with n≥ 5 per group from 2 independent experiments. Data were analyzed by Student’s t-test; *P < 0.05; **P < 0.01; ***P < 0.001; ****P < 0.0001.

We examined the frequency, total numbers, and phenotype of CD4^+^ T cells in the CNS of EAE mice that were i.v. tolerized with AST-EVs and treated with anti-IFN-γ mAb, and found significantly greater total numbers of IL-17A- and TNF-producing CD4^+^ T cells in mice treated with anti-IFN-γ mAb than those treated with isotype control mAb, but no significant differences were observed in their frequencies (**Fig. 9A**). The frequency and the absolute number of CD4⁺IL-10⁺ cells were not different in mice treated with anti-IFN-γ mAb compared to those treated with isotype control mAb (**Fig. 9B**). Treatment with anti-IFN-γ mAb reduced the percentage of PD-1^+^CD4^+^ T cells (**Fig. 9C**) and decreased apoptosis in CD4^+^ T cells. To further investigate AQP4_201-220-_specific CD4^+^ T cell response in anti-IFN-γ mAb-treated mice, we measured cytokine production in responses to AQP4_201–220_. Splenocytes of anti-IFN-γ mAb-treated mice secreted more GM-CSF, IL-17, and IFN-γ but lower IL-10 compared to those treated with isotype control mAb (**Fig. 9D**). These results demonstrate that AST-EVs suppress the disease in an IFN-γ-dependent manner.

**Fig. 9.**
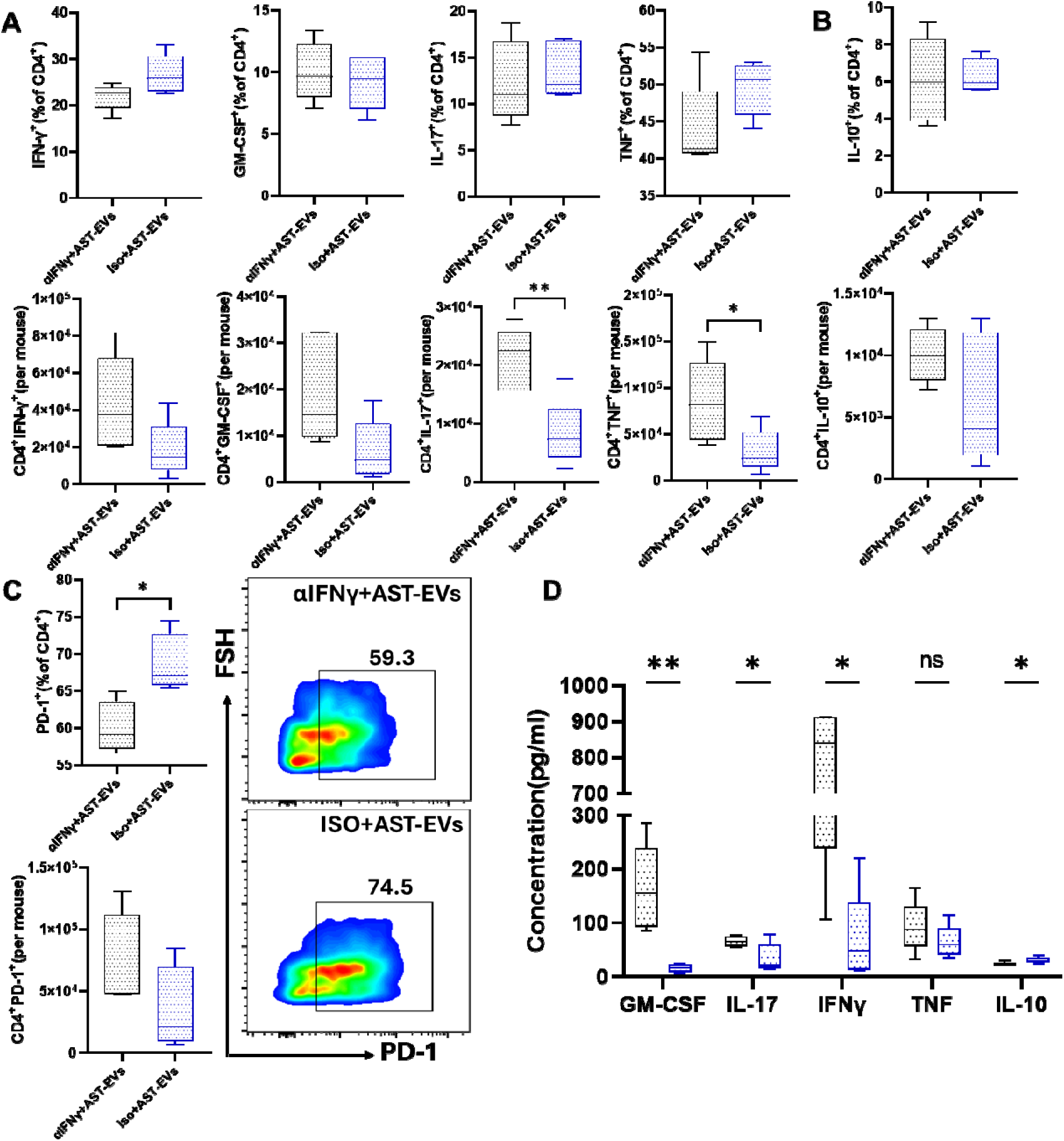
IFN-γ mediates cytokine production and PD-1 expression of Th cells in i.v. tolerance induced by AST-EVs. Mice shown in Fig. 8 were sacrificed on day 23 p.i., and MNCs isolated from the CNS and spleen were analyzed. (A). Frequencies and numbers of Th cells from the CNS of eNMOSD mice treated with AST-EVs and anti-IFN-γ mAb or isotype control mAb. (B) Frequencies of IL-10^+^CD4^+^ T cells from the CNS. (C) Representative flow cytometry plot of PD-1^+^ cells in gated CD4^+^ cells. (D) Cytokine concentrations in the supernatants of splenocyte cultures stimulated with AQP4 in mice treated with anti-IFN-γ mAb or Iso mAb. Data represent mean ± SEM (n=5 each group). Data were analyzed by Student’s t-test; *P < 0.05; **P < 0.01.

## Discussion

Existing therapies for NMOSD have been very beneficial in controlling disease progression, but remain largely Ag-nonspecific and immunosuppressive, exposing patients to serious, sometimes long-term risks. Consequently, prolonged use of these therapies correlated with complications of systemic immunosuppression (26, 27). Given that the pathogenesis of NMOSD is focused on a well-defined autoantigen (AQP4) in a significant proportion of patients, there is a compelling rationale to develop antigen-specific tolerance approaches that silence pathogenic responses while leaving the rest of the immune system intact.

Since NMOSD is a rare disorder with no reliable animal model, only a few studies have investigated tolerance induction in NMOSD using animal models. For instance, one study reported that the AQP4_201-220_ peptide dramatically decreased disease incidence and symptom severity; however, this model relies on the use of IFN-γR-deficient mice or anti-IFN-γ-treated mice (7). Tolerance induction, using peptide-loaded tolerogenic dendritic cells, has been tested in a phase Ib clinical trial in patients with MS, as well as in seropositive and seronegative NMOSD (28). Despite this considerable progress, translation to clinical application still requires overcoming several obstacles, including optimizing antigen delivery, ensuring long-term efficacy, and identifying reliable biomarkers (28, 29). Furthermore, more than 10 epitopes of AQP4 have been identified as pathogenic for AQP4-specific T cell responses, primarily targeting Th17 cells, in NMOSD patients (34–36), as well as multiple epitopes in animal models (7, 9, 16). While the majority (∼80%) of NMOSD patients have a pathogenic serum IgG against AQP4 (30), about 20-30% of patients are seronegative for AQP4 Ab. The target Ags most likely are other astrocytic Ags, AQP1, for example, or non-astrocytic Ags (31, 32). Overall, the identity of relevant Ag(s) in NMOSD remains unknown, with the possibility of heterogeneity among patients and over time. It is this lack of knowledge about Ags that hampers the development of Ag-specific NMOSD therapies, while using whole proteins risks activating non-targeted or even pathogenic immune responses (5). It is thus of great necessity to develop Ag-specific therapies targeting all AQP4 epitopes and other potential astrocytic antigens in NMOSD, despite the lack of exact epitope knowledge.

In recent years, EVs have garnered considerable interest due to their pivotal roles in intercellular communication and their therapeutic potential. EVs are bilayer-encapsulated vesicles released by nearly all cell types, carrying proteins, lipids, and nucleic acids that can modulate immune and neural responses under both physiological and pathological conditions (33). Previously, we demonstrated that treatment with oligodendrocyte-derived EVs (OL-EVs), administered either before or after disease onset, reduced the severity of neurological disease and CNS inflammation in EAE mice. The therapeutic properties of OL-EVs in EAE were shown to depend on the presence of myelin Ags in EVs derived from OL cultures (11). Limited research has been conducted on how EV production and how astrocytes, a key cellular target in NMOSD, change during the disease, as well as their therapeutic potential in suppressing disease progression (34, 35). The goal of this study was to induce tolerance in eNMOSD mouse model through AST-EVs. In the present study, for the first time, we demonstrate that AST-EVs treatment after disease onset significantly reduces disease severity, with inhibited AQP4201–220 peptide-induced, but not anti-CD3/anti-CD28-induced T cell proliferation, indicating that i.v. injection of AST-EVs suppressed eNMOSD in an AQP4-specific manner without affecting non-specific immune responses.

Many preclinical studies (notably work with mesenchymal and neural-lineage exosomes such as Ol-EVs) have reported that EVs can suppress neuroinflammation by reprogramming microglia/macrophages, eliciting tolerogenic antigen-presenting cells, inducing regulatory T cells, and increasing anti-inflammatory cytokines such as IL-10 (36). Mechanistically, CNS-derived EVs are preferentially taken up by phagocytes and monocytes. For instance, Ol-EVs have been shown to restore antigen-specific tolerance by inducing immunoregulatory monocytes that up-regulate PD-L1 and IL-10, and by promoting apoptosis of autoreactive CD4⁺ T cells, effects that were reported to be safer than infusion of free peptides in experimental models (11). It has been determined that IL-10, as an anti-inflammatory cytokine with a pivotal role in immune regulation, suppresses inflammatory responses in NMOSD (37) and is required for peptide/i.v. tolerance induction in NMOSD (28). Given that AQP4-specific CD4^+^ T cells play a pivotal role in B cell activation and antibody production during the NMOSD disorder progression (7, 38), decreased infiltrated AQP4-specific CD4^+^ T cells into the CNS by AST-EVs can be correlated with the suppression of CNS inflammation in eNMOSD. Furthermore, our study demonstrates that AST-EV treatment results in increased IL-10 production in CD4⁺ T cells, accompanied by enhanced PD-1 and FoxP3 expression, but decreased production of proinflammatory cytokines IL-17 and GM-CSF, indicating that AST-EV treatment promotes an anti-inflammatory immune response.

The role of B cells in NMOSD pathogenesis has been demonstrated through their ability to differentiate into antibody-producing plasma cells and facilitate Th17 differentiation by functioning as APCs and by producing inflammatory cytokines (39). In addition, because of the vital role of B cells in NMOSD, anti–CD20 treatment is now a common therapy for patients (40). Our study showed that AST-EV therapy significantly reduced the percentage of B cells in the CNS, which is consistent with previous findings and supports a strong association between CNS B-cell burden and NMOSD pathology (7). Specifically, autoimmune-associated B cells (ABCs; defined as CD19^+^CD11c^+^) are a subset of potent APCs in autoimmune diseases, expressing high levels of CD80/CD86 and MHC class II. These cells are believed to be a key feature in certain types of autoimmunity and are often increased in systemic lupus erythematosus, rheumatoid arthritis, and MS (41, 42). CD11c⁺ B cells are also significantly increased in patients with NMOSD, and this increase is associated with brain atrophy and disease severity, indicating that these cells contribute to neuroinflammation (43). Owing to these pathogenic properties, CD11c⁺ B cells are considered a potential target for immunotherapeutic intervention (44). We show that, upon treatment of eNMOSD mice with AST-EVs, the CD11c⁺ B cell population in the CNS is reduced. On the other hand, lower levels of IL-12p35 or IL-35 are strongly correlated with the severity of NMOSD (45, 46). IL-35 also prevented neuropathy in EAE mice by inhibiting the proliferation of pathogenic Th17 and Th1 cells and blocking the infiltration of inflammatory cells into the CNS (47). In our study, we found an increased proportion of IL-35^+^CD19^+^ regulatory B cells (Bregs) (23) in the CNS of eNMOSD mice after AST-EV treatment. This, together with the reduced proportion of ABCs and total B cells, demonstrates that the induction of tolerogenic B cells could be a novel mechanism underlying AST-EV-induced i.v. tolerance in eNMOSD.

The critical role of IFN-γ in modulating CNS autoimmunity, particularly through regulating the balance between disease-associated Th1 and Th17 responses, has been well established. (7, 48). We have previously demonstrated that i.v. injection of auto-Ag in EAE mice results in elevated production of IFN-γ, and blockade of IFN-γ reduces IL-27 production and PD-L1 expression in CNS monocyte-derived DCs, thereby abrogating tolerance induction (20, 25). Consistent with these reports, we demonstrate here that i.v. administration of anti–IFN-γ mAb prior to AST-EVs injection in mice with eNMOSD failed to suppress disease progression. The critical role of IFN-γ as a master regulator of immunopathogenesis of NMOSD has been shown by findings that IFN-γR-deficient mice or mice receiving neutralizing anti-IFN-γ mAb develop notably more severe eNMOSD than WT mice (7). However, immune tolerization could be induced in this model with AQP4_201-220-_-coupled poly(lactic-co-glycolic acid) nanoparticles even after IFN-γ blockade, suggesting that IFN-γ signaling may be important for counteracting disease induction but not for tolerance induction. The precise mechanisms involved in this pathway remain to be elucidated.

In summary, our results demonstrate for the first time that AST-EVs can exert a therapeutic effect by competently attenuating disease progression in mice with eNMOSD. Given that AST-EVs contain a spectrum of astrocytic Ags, they can potentially induce Ag-specific tolerance in NMOSD and other astrocyte-related neurological diseases. Furthermore, EV-based therapeutics have been reported to be well-tolerated in both preclinical and clinical contexts (49). Together, AST-EVs could provide a safe and disease-specific approach to restoring immune homeostasis while avoiding the risks of generalized immunosuppression, thus representing a promising novel Ag-specific treatment strategy for NMOSD.

## Acknowledgements

The author(s) declare that financial support was received for the research and/or publication of this article. This work was supported by R01 AI152251 and R01 AI160189 grants from the National Institutes of Health (NIH) to Abdolmohamad Rostami.

## Disclosures

The authors declare that no competing interests exist.

## Notes

### Competing Interest Statement

The authors have declared no competing interest.

