## Supplemental Figures for "Astrocyte-derived extracellular vesicles as antigen-specific therapy for neuromyelitis optica spectrum disorder in the mouse model"

**Supplementary Materials**

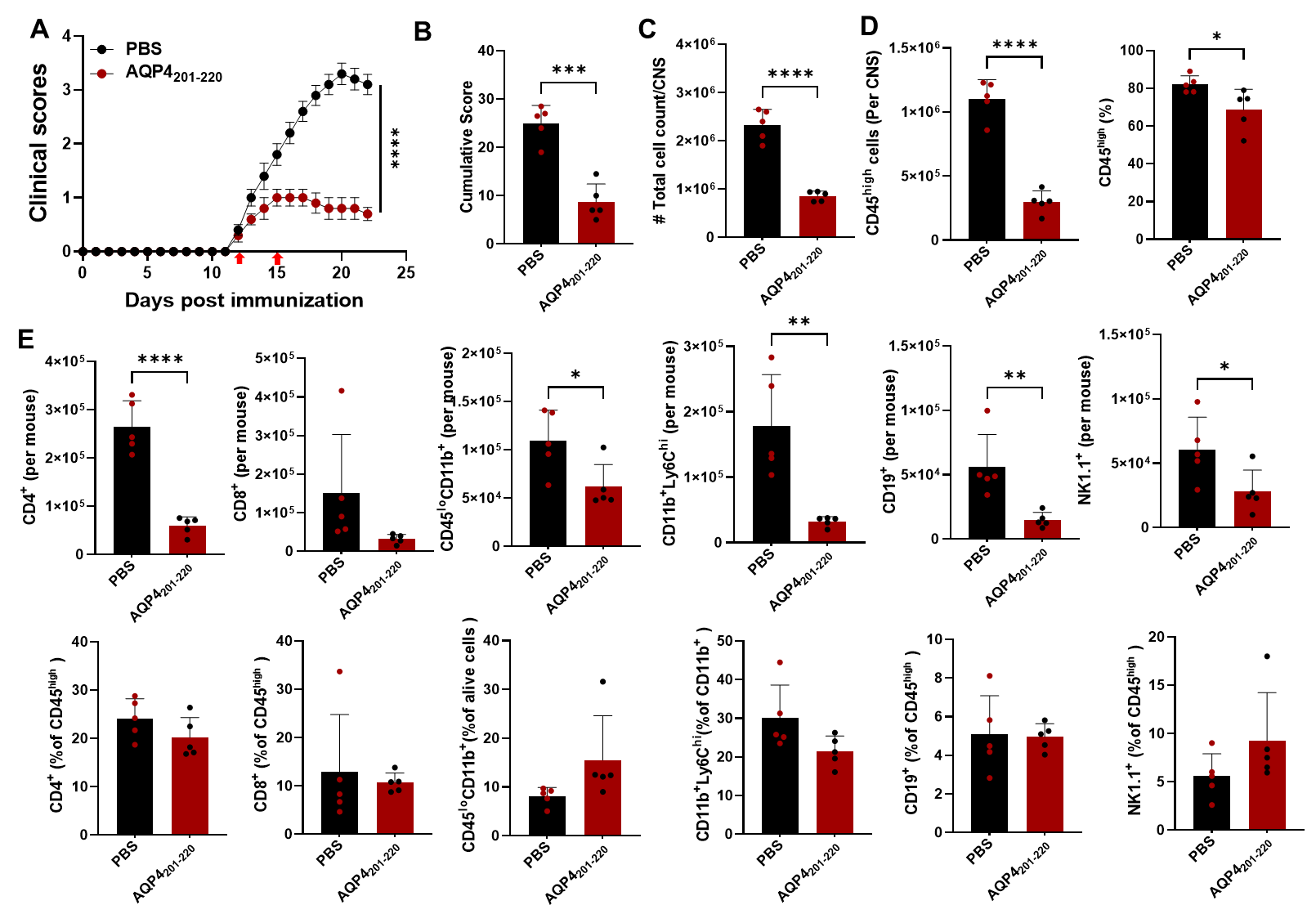

**Fig. S1.** **AQP4_201-220_ i.v. injection suppressed NMOSD progression in the mouse model.** (A) Mice with eNMOSD (n = 5 per group in two experiments) were treated via i.v. injection with AQP4_201-220_ peptide (50 μg per mouse per injection) or PBS on days 12 and 15 post-immunization (p.i.). (B) Cumulative disease severity scores. (C) Mice were sacrificed on day 23 p.i., the spinal cord was harvested after extensive perfusion, and the total number of cells infiltrating the CNS was counted. (D) The numbers of CD45^high^ leukocytes in the CNS were determined by flow cytometry. (E) The absolute numbers and frequencies of CD4^+^ T cells, CD8^+^ T cells, CD45^low^CD11b^+^ (microglia), CD11b^+^Ly6C^high^ (monocytes), B cells, and NK cells are also evaluated. Data shown as mean ± SEM. Values are mean ± SD. Unpaired t-test; p < 0.05 (*), p < 0.01 (**), p < 0.001 (***), and p < 0.0001 (****) were considered significant.

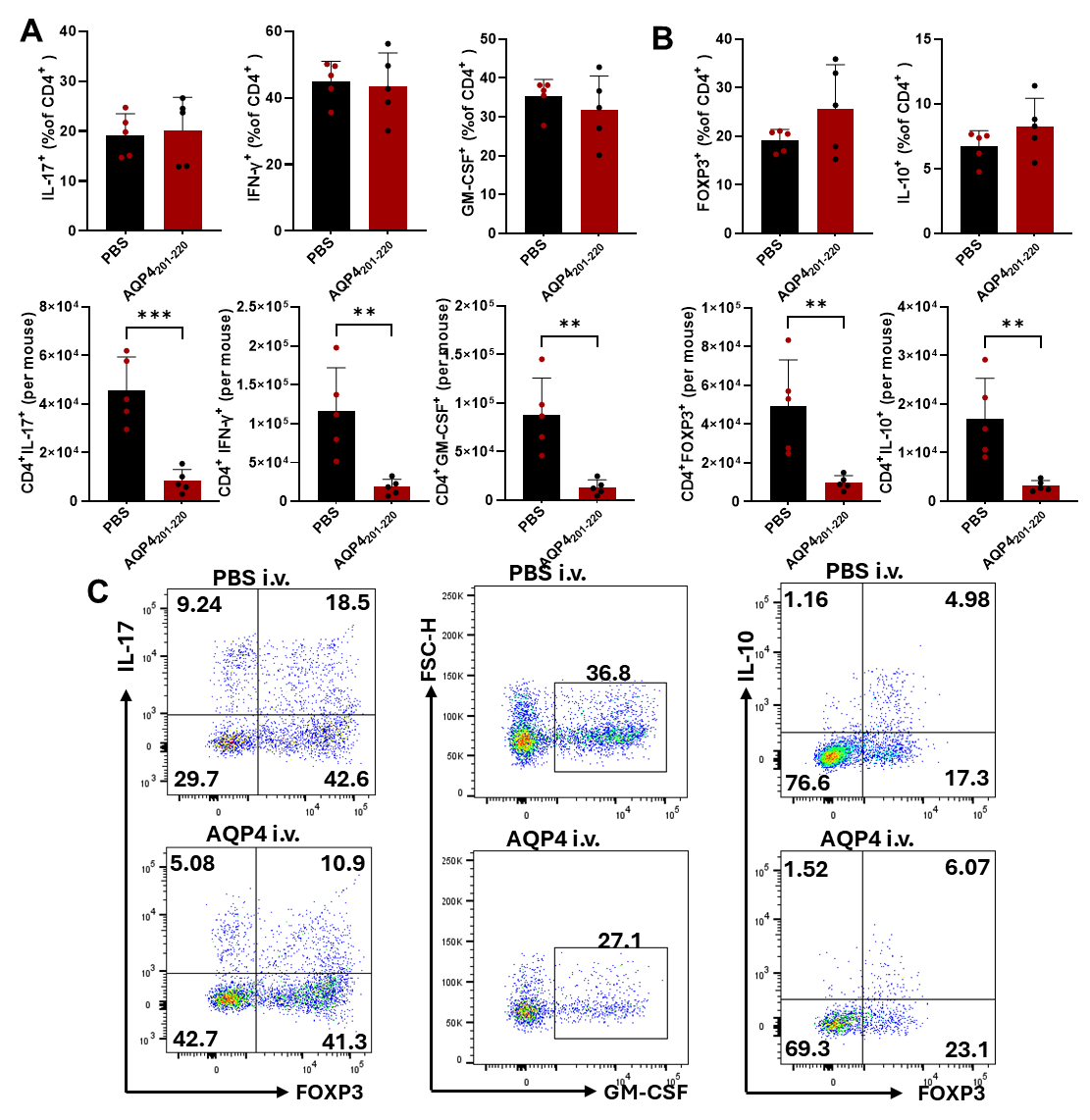
.

**Fig. S2. Frequency and absolute number of cytokine-producing T cells in the** AQP4_201-220-_ **and PBS-treated mice.** (**A**). Higher absolute numbers of cytokine-producing CD4^+^ T cells were observed in the PBS-treated group during eNMOSD. (**B**) IL-10^+^ and FOXP3^+^ CD4^+^ T cell frequencies were slightly higher in AQP4_201-220_-treated mice. (**C**) Representative flowcytometry plot of IL-17A^+^, IFN-γ^+^, GM-CSF^+^, FOXP3^+^, and IL-10^+^ cells in gated CD4^+^ cells of both groups. Values are mean ± SD (n=5 each group). Unpaired t-test; ** p < 0.01, *** p < 0.001.

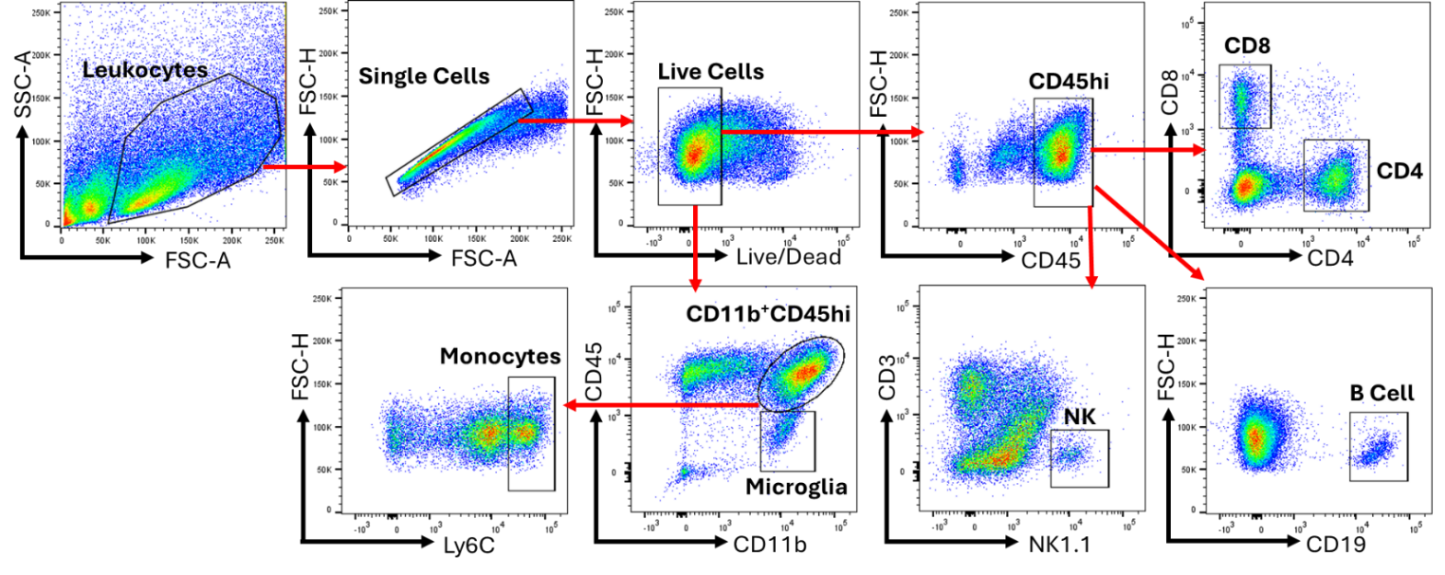

**Fig. S3. Gating strategy for immune cell subsets of the CNS.** Gating strategy for flow cytometric analysis of lymphoid and myeloid cells in the CNS of eNMOSD mice. A similar gating strategy was used in analyses of immune cells from other organs. Representative plots of gating strategy for immune populations in the CNS.
